# Delineating the relationship between immune system aging and myogenesis in muscle repair

**DOI:** 10.1101/2020.03.23.003160

**Authors:** Stephanie W. Tobin, Faisal J. Alibhai, Lukasz Wlodarek, Azadeh Yeganeh, Sean Millar, Jun Wu, Shu-hong Li, Richard D. Weisel, Ren-Ke Li

**Affiliations:** Toronto General Research Institute, Division of Cardiovascular Surgery, University Health Network, Toronto, Canada

**Keywords:** Aging, Inflammation, Bone Marrow, Myogenesis

## Abstract

How aging affects the communication between immune cells and myoblasts during myogenesis is unclear. We therefore investigated how aging impacts the cellular synchronization of these two processes after muscle injury. Muscles of old mice (20 months) had chronic inflammation and fewer satellite cells compared to young mice (3 months). After injury, young mice developed a robust, but transient inflammatory response and a stepwise myogenic gene expression program. These responses were impaired with age. Replacement of old bone marrow (BM) via heterochronic bone marrow transplantation (BMT) increased muscle mass and performance on locomotive and behavioural tests. After injury, Y-O BMT restored the immune cell and cytokine profiles to a young phenotype and enhanced satellite cell activity while O-O BMT amplified a late-onset proinflammatory response. *In vitro,* conditioned media from young or old macrophages had no effect or impaired myoblast proliferation, respectively. Thus, BM age negatively affects myogenesis by inhibiting myoblast proliferation.

## Introduction

Skeletal muscle contains tissue-resident muscle stem cells called satellite cells which can proliferate and differentiate into mature muscle. The satellite cell population is limited, however, as extensive activation caused by chronic injury may exhaust the stem cell pool ^1^. Aging also results in homeostatic and regenerative defects that leads to muscle loss via signals that are both intrinsic and extrinsic to the satellite cell ^2^. These signals ultimately impact absolute progenitor cell number and function.

Satellite cells become inherently defective with age, as transplantation of old satellite cells to young muscle fail to repair muscle effectively ^3,4^. Intrinsic factors that may account for satellite cell aging include increased DNA damage, increased expression of cell cycle checkpoint regulators such as p16 and p53, and loss of H3K27me3 histone methylation via reduced Ezh2 activity ^5–7^. Aged muscle is subject to higher levels of muscle turnover by changes in the satellite cell niche which may exhaust the stem cell pool ^8^. With age, satellite cells also become prone to higher levels of symmetric division, leading to fewer Pax7^+^ progenitor cells ^4^, and geriatric satellite cells may also enter an irreversible state of quiescence ^7^. Myogenesis may also be influenced by extrinsic factors in the blood that differ with age. This has been documented using heterochronic parabiosis and blood transfusion ^9–11^. Circulating factors, particularly members of the TGF-β superfamily, increase with age and contribute to muscle wasting via upregulation of cyclin dependent kinase inhibitors ^12^. Despite moderate improvements in aged mice that receive young blood ^11^, the aged muscle microenvironment limits regenerative capacity.

In addition to satellite cells, there are muscle-resident cells such as macrophages, endothelial cells, Hoechst dye-excluding Side Population cells, and mesenchymal stem cells. Some of these cells arise from circulating cells which originate from the bone marrow. These cells alter the satellite cell microenvironment and modulate satellite cell maintenance. They are largely distinguished by their cell surface markers: CD45^+^, CD31^+^ and Sca-1^+^/CD45^−^/CD31^−^ are generally associated with hematopoietic, endothelial and mesenchymal stem cells, respectively. Under normal conditions, muscle damage induces the robust recruitment of blood-derived inflammatory cells including monocytes, to complete muscle repair ^13–15^. Macrophage depletion alters satellite cell proliferation, prevents myogenesis and results in fibrosis ^13–15^. Aged muscle suffers from macrophage deficiencies which lead to chronic inflammation, fibrosis ^16^ and a delayed inflammatory and myogenic response after injury ^17,18^. Recently, Wang et al. showed that sarcopenia could be delayed when aged mice received bone marrow transplant with young donor cells ^19^ but the populations, timing and factors released by various inflammatory cells has not been adequately characterized.

Based on these studies, we sought to understand how bone marrow aging affects the satellite cell population during rest and after acute injury in young vs aged mice or after Bone Marrow Transplant of young or old Sca-1^+^ cells into aged mice. Using comprehensive flow cytometry analysis we profiled changes in myogenic bone marrow derived cells in injured muscle across all groups and profiled cytokine and muscle gene expression patterns. We hypothesized that young bone marrow could rescue satellite cell function after acute muscle injury through differential recruitment of inflammatory cells and their respective cytokines.

## Results

### The aged muscle microenvironment is enriched in neutrophils and T cells

We first characterized the physiological differences of skeletal muscles between young (3 month old) and old (20 month old) mice. With age, the weight of the *Tibialis anterior* (TA) muscle decreased, as expected (Figure 1A-B). There was no difference in the frequency distributions of myofibers between these age groups, although the average cross-sectional area trended lower in older mice (Figure 1C-E). We then performed three locomotive and behavioural tests: using the open field test we assessed ambulatory distance and rearing count and using the rotarod we measured the average time to fall. In all cases, old mice did significantly worse than young mice which are an indirect readout of indicative of impaired muscle function (Figure 1F-H).

**Figure 1.**
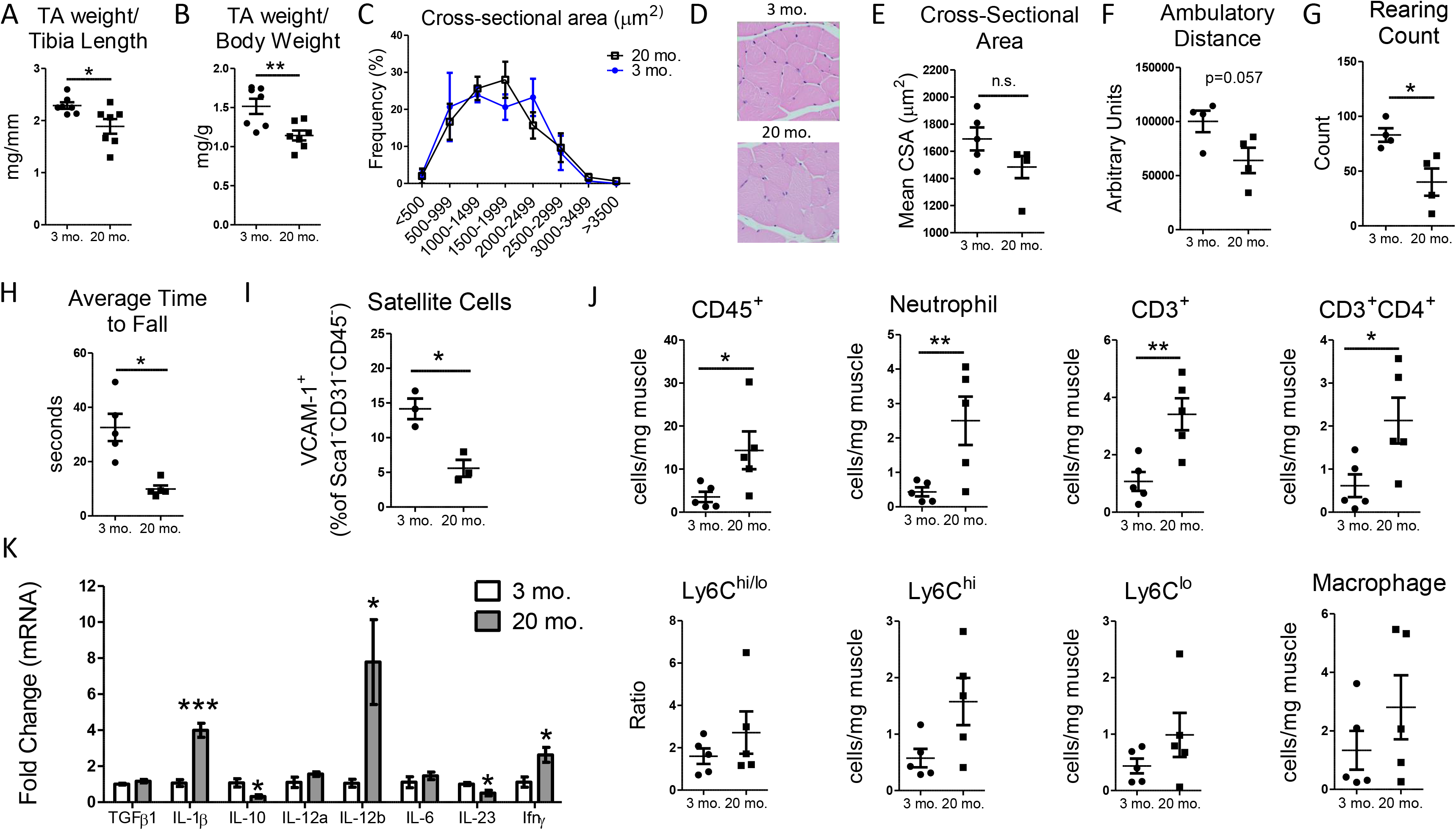
Age-associated muscle deficiencies are coupled with inflammation. A, B) TA weight (normalized to tibia length or body weight) is reduced with age (n=7). For all experiments in this figure, young mice were 3 months old and old mice were >20 months old. C) Frequency plot of the CSA of the TA muscle from 3- and 20-month old mice grouped into 500 μm^2^ bins (n=5). D) Representative image of H&E staining of TA histological sections of 3- and 20-month old mice. Scale = 50 μm. E) Average CSA (μm^2^) of the TA muscle from 3- and 20-month old mice (n=5). F-G) The open field test was used to assess ambulatory distance (F) and rearing count (G) (n=4). H) A rotating beam (rotarod) was used to quantify the average time to fall (n=5). I-J) Flow cytometry analysis of satellite cells (n=3) or inflammatory cells (n=5) and corresponding flow panels. Satellite cells are plotted as %VCAM-1^+^ of the Sca-1^−^, CD45^−^ and CD31^−^ population. Immune cells are plotted as cells/mg muscle. K) Expression analysis of cytokines in the gastrocnemius muscles of young and old mice (n=4). The data are reported as relative to housekeeping gene, β-actin. All data are presented as mean±SEM. All analyses were done using an unpaired t-test (2 groups) or two-way ANOVA (≥2 factors). *P<0.05, **P<0.01, ***P<0.001. TA, Tibialis Anterior; CSA, Cross-Sectional Area. H&E: Hematoxylin and eosin.

We next focused on profiling the cellular differences between young and old muscles in terms of satellite cells, myeloid cells and T cells. Aging negatively impacted the satellite cell population (Figure 1I) while CD45^+^ cells inflammatory cells were increased (Figure 1J). CD45^+^ cells were primarily neutrophils and CD3^+^/CD4^+^ T cells. The number of Ly6C^hi^ and Ly6C^lo^ monocytes and F4/80^+^ macrophages was not affected by age. In terms of cytokine expression, aging increased the expression of IL-1β, IL-12b, and IFNγ but reduced expression of IL-23 and IL-10 in the Gastrocnemius muscle (Figure 1K).

### Aged muscle has a damaged immune response after injury

To investigate how age affects muscle repair, we profiled cytokine and cellular infiltration of immune cells to muscle after acute injury by Cardiotoxin (CTX). We profiled the expression of six cytokines in Control, 3- and 10-days post CTX in the TA muscle. Genes were grouped into M1 or M2 categories for simplicity. Pro-inflammatory cytokines, such as IL-1β and IL-12b were elevated in old TA muscle, as observed in Gastrocnemius muscle (Figure 2A and Figure 1K). In young muscle, the general trend of IL-1β and IL-12b was upregulation at 3 days post CTX and a return to baseline levels by 10 days post CTX. IL-12a and IL-23 tended to be lower in old mice at 3- and 10-days post CTX. IL-10 was significantly higher in old muscle at 3 days post injury compared to young muscle. We examined iNOS and Arg1 expression which canonically denote inflammatory M1 and anti-inflammatory M2 macrophages, respectively. iNOS1 was higher in naïve muscle of Old mice while Arg1 expression increased 3 days post CTX in young but not old muscle.

**Figure 2.**
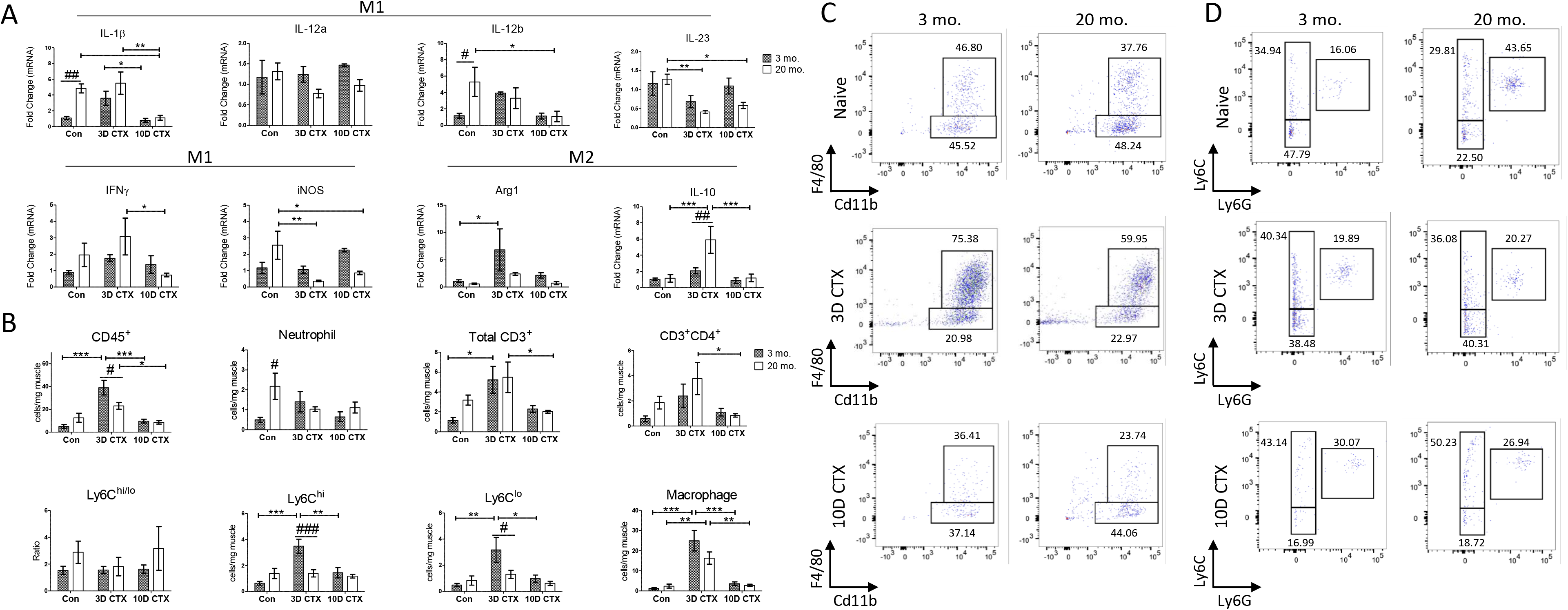
Young muscle produces a robust and transient inflammatory response that is impaired with age. A) Gene expression analysis of cytokines broadly clustered as M1 or M2 markers in the TA muscle of young (3 month old) and old (20 month old) mice 3- and 10-days post CTX injury. The control (Con) group is TA muscle that received an injection of saline. The qRT-PCR data are reported as relative to housekeeping gene, Hprt (n=4). B) Flow cytometry analysis of immune cells in naïve muscle and at 3- and 10-days post CTX injury. Naïve flow cytometry data is derived from Figure 1K. Naïve n=6, 3D CTX n=7, 10D CTX n=5. Immune cells are plotted as cells/mg muscle. C) Flow cytometry gating of macrophages in naïve muscle and at 3- and 10-days post CTX injury (CD45^+^CD11b^+^F4/80^+^). D) Flow cytometry gating of Ly6C^hi^ and Ly6C^lo^ monocytes (Ly6C^+^Ly6G^−^) and neutrophils (Ly6C^+^Ly6G^+^). Percentages of different cell populations are shown in the indicated box. All data are presented as mean±SEM. All analyses were done using an unpaired t-test (2 groups) or two-way ANOVA (≥2 factors). ***P<0.05, **P<0.01, ***P<0.001 delineate differences within young or old cohorts. #P<0.05 delineate differences between young and old cohorts. TA, Tibialis Anterior; CTX, Cardiotoxin; Con, Saline Control.

We next assessed the recruitment of neutrophils, macrophages, monocytes and T cells. In 3 month old mice, overall CD45^+^ cells were significantly higher at 3 days post injury, and returned to near normal numbers by 10 days post injury. Although older muscle showed a similar trend, the infiltration of CD45^+^ at 3 days post injury was significantly lower than in young muscle (Figure 2B). Based on macrophage number normalized to muscle weight (Figure 2A), this cell population accounts for the majority of infiltrating CD45 cells at 3 days post CTX in both ages. As exemplified in representative flow cytometry images, macrophages rapidly infiltrate injured muscle of young and old mice at 3 days post CTX and largely disappear by 10 days post CTX (Figure 2C). Compared to naïve muscle, old mice had a reduction in neutrophils at 3- and 10-days post injury and no monocyte recruitment (Figure 2B and representative flow cytometry images in Figure 2D. At three days post CTX, the total CD3^+^ T cell population was not affected by age, by the CD3^+^CD4^+^ T cell population trended higher in old mice. Thus, although old muscle has higher pro-inflammatory signals at baseline, there is an inefficient inflammatory response after injury.

### Heterochronic bone marrow transplant did not alter the immune populations in naïve skeletal muscle

To determine the extent to which inflammatory cell age affects myogenesis in old muscle, we used heterochronic bone marrow transplant. Sca-1^+^ enriched bone marrow (BM) cells from young or old GFP^+^ donor mice were transplanted into lethally irradiated aged recipients as previously described ^20^. Three months after BM transplant, hindlimb muscle was collected for comprehensive flow cytometry, gene expression and locomotive analysis. When reconstituted with young Sca-1^+^ BM (Y-O), the percentage of GFP^+^ cells present in muscle was equal to mice that received old Sca-1^+^ BM (O-O) (Figure 3A). Greater than 90% of all GFP^+^ mononuclear cells in muscle were CD45^+^, irrespective of donor age (data not shown). TA muscle mass was higher in Y-O mice when normalized to tibia length but not body weight (Figure 3B-C). No significant differences in average cross-sectional area or frequency distributions were observed (Figure 3D-F). With respect to behavioural and locomotive tests, young bone marrow improved ambulatory distance, rearing count and the average time to fall (Figure 3G-I). We found no difference in the frequency of satellite cells in either group, all of which were GFP^−^ (Figure 3J). The bone marrow contributed to a low frequency of GFP^+^MSCs (less than 0.5% of mononuclear cells) (Figure 3K). Finally, donor bone marrow age did not affect the number of GFP^+^ immune cells present in muscle (Supplemental Figure S1). Thus, although we identified improved muscle mass and locomotive and behavioural outputs in old mice that received young Sca-1^+^ bone marrow cells, we could not attribute this to changes in satellite cell, MSC or immune cell populations.

**Figure 3.**
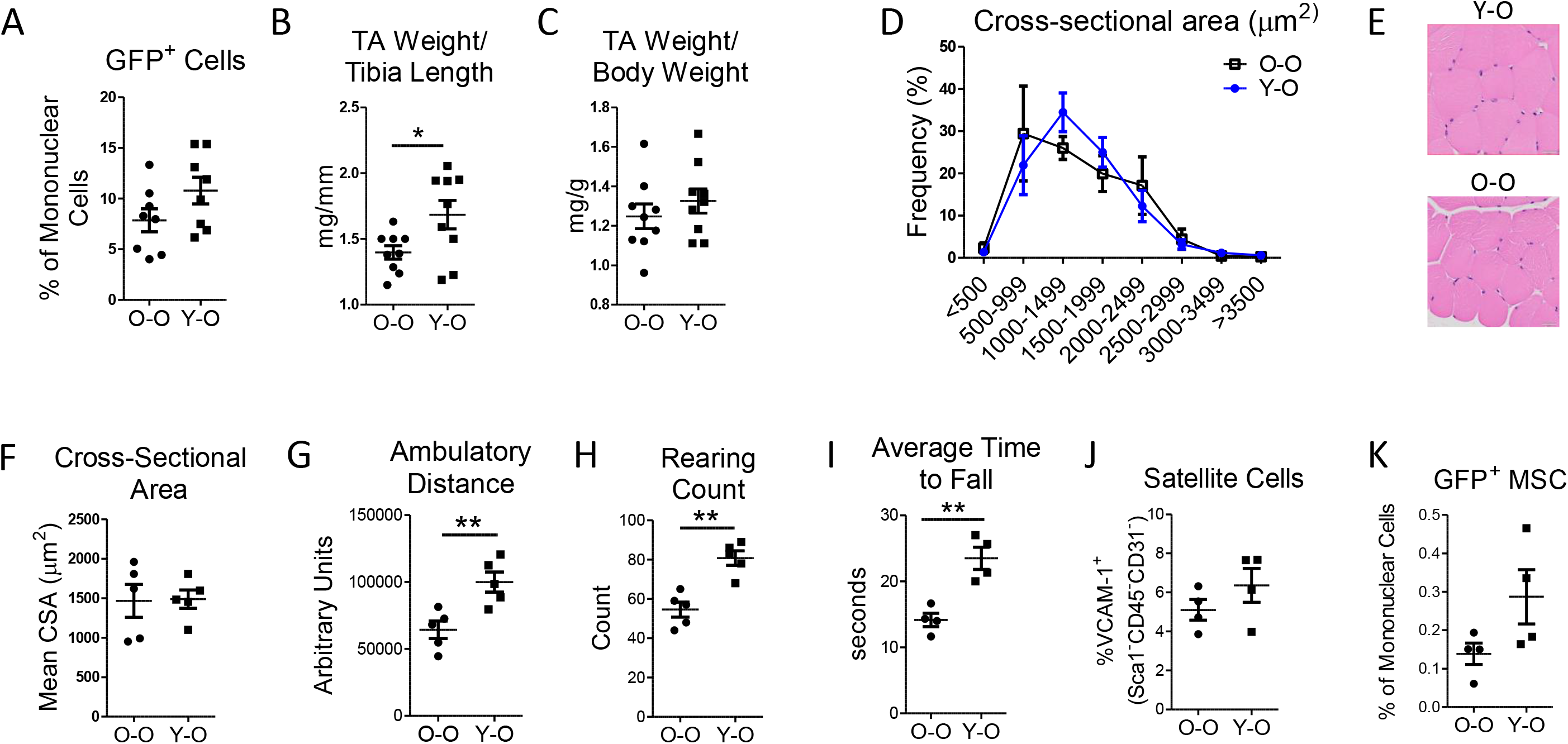
Muscle function and cellular profiles in aged muscle after heterochronic bone marrow transplant. A) Percentage of GFP^+^ cells in hindlimb muscle after heterochronic Bone Marrow Transplant of Sca-1^+^GFP^+^ cells (n=8). Mice that received young Sca-1^+^ BM cells are labelled Y-O. Mice that received old Sca-1^+^ BM cells are labelled O-O. All recipients (hosts) were >18 months old at the time of reconstitution and underwent a 12-week recovery period. B-C) TA weight (normalized to tibia length or body weight) (n=9). D) Representative image of H&E staining of TA histological sections of Y-O and O-O mice. Scale = 100 μm. E) Average CSA (μm^2^) of the TA muscle from Y-O and O-O mice (n=5). F) Frequency plot of the CSA of the TA muscle from Y-O and O-O mice grouped into 500 μm^2^ bins (n=5). G-H) The open field test was used to assess ambulatory distance (G) and rearing count (H) (n=5). I) A rotating beam (rotarod) was used to quantify the average time to fall (n=4). J-K) Flow cytometry analysis of GFP^−^ satellite cells and GFP^+^ mesenchymal stem cells (n=4). Satellite cells are plotted as %VCAM-1^+^ of the Sca-1^−^, CD45^−^ and CD31^−^ population; MSCs are plotted as % Sca-1^+^, CD45^−^ and CD31^−^ of mononuclear cells. All analyses were done using an unpaired t-test (2 groups) or two-way ANOVA (≥2 factors). *P<0.05, **P<0.01. TA, Tibialis Anterior; CSA, Cross-Sectional Area; MSC, Mesenchymal Stem Cell.

### Bone marrow aging alters leukocyte recruitment to injured muscle

To determine the age-related role of bone marrow-derived cells in the muscle inflammatory response of muscle repair, reconstituted mice were injected with Cardiotoxin (CTX) and the percentage of GFP^+^ cells was assessed by flow cytometry 3- and 10-days later. Mononuclear cells in injured skeletal muscle were composed of 50-70% GFP^+^ cells (Figure 4A). This is dramatically higher than the 5-10% of GFP^+^ cells observed in naïve muscle. Interestingly, O-O recipients had a higher proportion of GFP^+^ cells at 10 days post injury. The total number of GFP^+^CD45^+^ cells normalized to muscle weight demonstrates the severity that age has on chronic immune cell infiltration to muscle: GFP^+^CD45^+^ cells peaked at 3 days post CTX and returned to baseline levels in Y-O, while in O-O this population increased further at 10 days post CTX (Figure 4B). Representative flow cytometry images of GFP^+^CD45^+^ cells show that while similar levels of GFP^+^CD45^+^ cells are present at 3 days post CTX, the O-O BMT cohort is chronically inflamed at 10 days post CTX (Figure 4C). Figure 4D depicts immunofluorescent staining of the GFP^+^CD45^+^ population between the Y-O and O-O groups at 3- and 10-days post injury. At 3 days post injury, GFP^+^CD45^+^ cells are present within the site of muscle damage. At 10 days post injury there is a higher amount of GFP^+^CD45^+^ in mice that received old BM.

**Figure 4.**
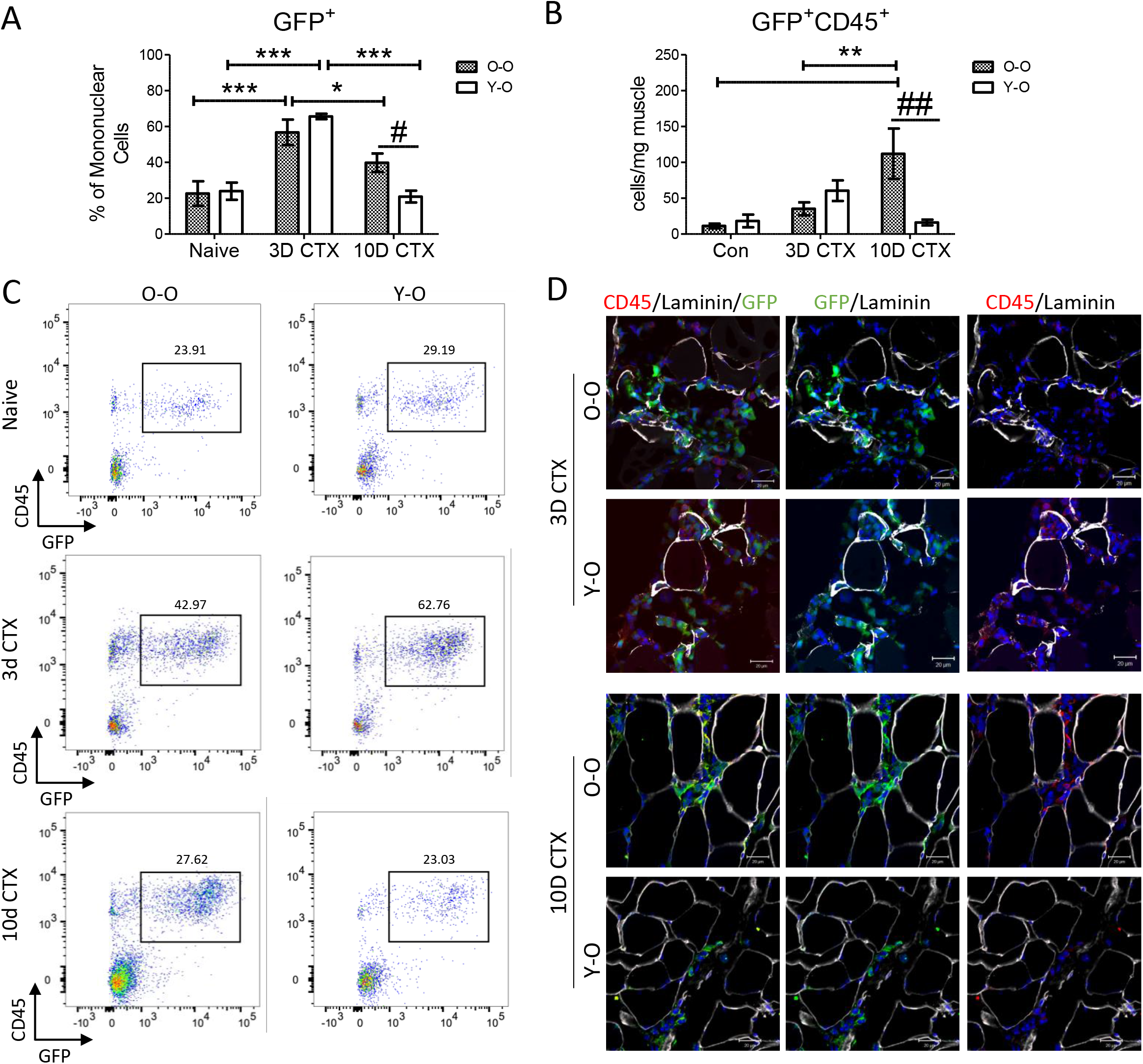
GFP^+^CD45^+^ cell infiltration resolves faster after heterochronic BMT in injured muscle. A) Percentage of GFP^+^ cells in hindlimb muscle after heterochronic bone marrow transplant of Sca-1^+^GFP^+^ cells and acute muscle injury using CTX. All recipients (hosts) were >18 months old at the time of reconstitution and underwent a 12-week recovery period before CTX administration. Mice that received young Sca-1^+^ BM cells are labelled Y-O. Mice that received old Sca-1^+^ BM cells are labelled O-O. Muscles were assessed 3- and 10-days post CTX injection. B) Number of GFP^+^CD45^+^ per mg muscle. Naïve n=4, 3D CTX n=5 10D CTX n=5. C) Flow cytometry gating of GFP^+^CD45^+^ cells. Percentages of the GFP^+^CD45^+^ cell populations are shown in the indicated box. D) Representative images of staining for Laminin (white), GFP (green) and CD45 (red) of TA cryosections of Y-O and O-O mice at 3- and 10-days post CTX. All data are presented as mean±SEM. All analyses were done using a two-way ANOVA (≥2 factors). **P<0.05, **P<0.01, ***P<0.001 delineate differences within Y-Y or Y-O cohorts. #P<0.05 delineate differences between Y-O and O-O cohorts. TA, Tibialis Anterior; CTX, Cardiotoxin; BMT, Bone Marrow Transplant.

### Bone marrow rejuvenation influences the myeloid and satellite cell populations of old mice

We completed gene expression analysis and flow cytometry for neutrophils, monocytes, macrophages and T cells at each time point to better understand which cells were responsible for the age-dependent differences we observed in CD45^+^ cell recruitment to injured muscle in old hosts. We first evaluated the expression of six cytokines during this repair process: IL-1β, IL-12a, IL-12b, IL-23, IFNγ and IL-10 (Figure 5A). Y-O had higher expression of IL-1β and IFNγ expression at 3 days post CTX compared to O-O. IL-10 followed a similar pattern. Notably, IL-1β expression increased in O-O much later at 10 days post CTX. IL-12b expression increased in both Y-O and O-O at 10 days post CTX as was its co-factor, IL12a, in O-O mice. In terms of cellular infiltration, we saw that monocytes were the main population affected by BM age as at 3 days post CTX there were more GFP^+^Ly6C^hi^ monocytes in Y-O recipient muscle (Figure 5B). GFP^+^ macrophages followed a similar trend, though based on cell staining for Mac-3 and flow cytometry we did not see age-related difference in overall macrophage infiltration (Figure 5B-D). The late onset of immune cell infiltration in O-O recipients at 10 days post CTX was derived from GFP^+^Ly6C^lo^ monocytes and GFP^+^T cells. GFP^+^ neutrophils were also more abundant at 10 days post CTX compared to Y-O (Figure 5B). Representative flow images show the recruitment of GFP^+^ monocytes and neutrophils (Figure 5E).

**Figure 5.**
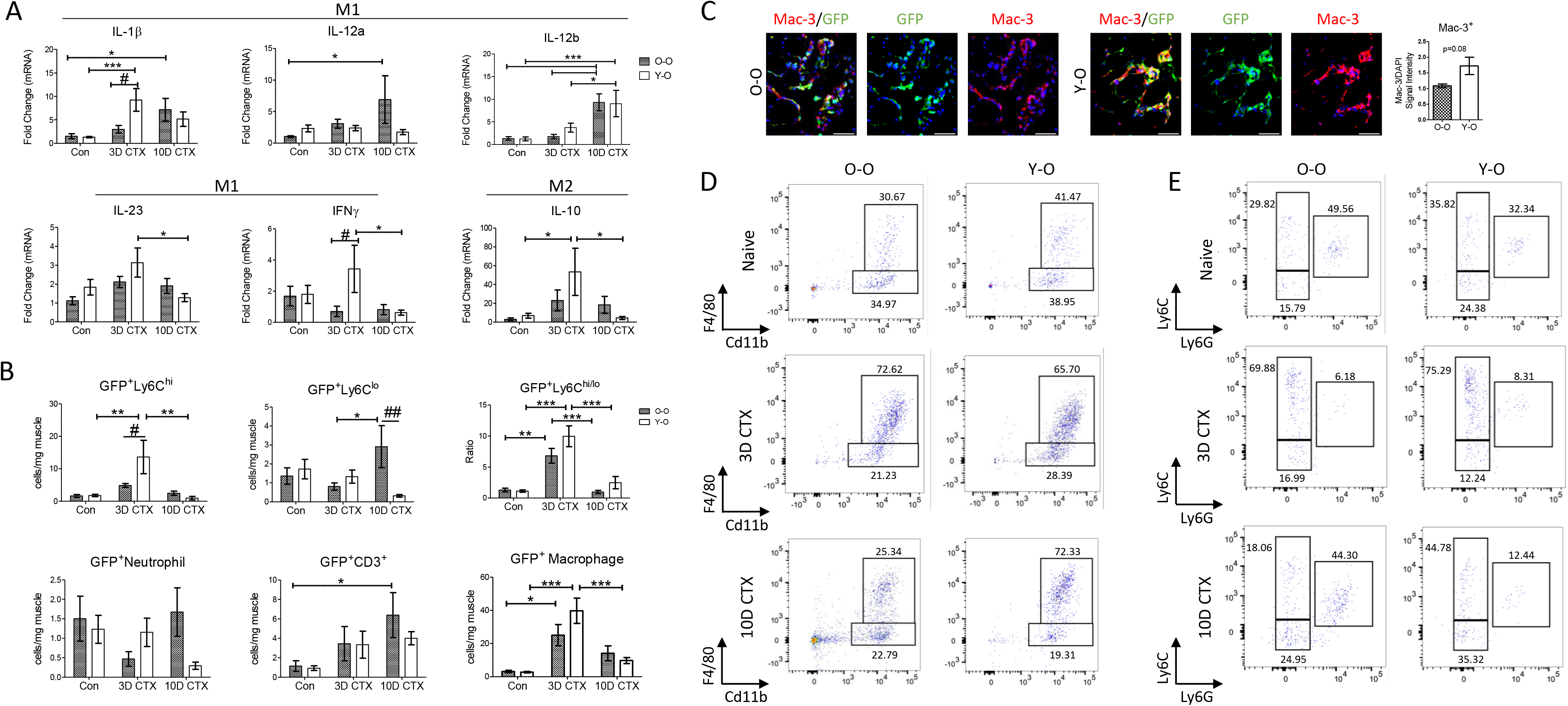
Young Sca-1^+^ BM cells restore the timing of the myeloid response after acute muscle injury in old mice. A) Gene expression analysis of cytokines in the injured TA muscle at 3- and 10-days post injury after heterochronic BMT. All recipients (hosts) were >18 months old at the time of reconstitution and underwent a 12-week recovery period before CTX administration. Mice that received young Sca-1^+^ BM cells are labelled Y-O. Mice that received old Sca-1^+^ BM cells are labelled O-O. The control (Con) group is TA muscle that received an injection of saline. The qRT-PCR data are reported as relative to housekeeping gene, Hprt (n=5-7). B) Flow cytometry analysis of GFP^+^ immune cells in injured muscle. Naïve n=4, 3D CTX n=5, 10D CTX n=5. Inflammatory cells are plotted as cells/mg muscle. Naïve flow cytometry data is derived from Supplementary Figure S1. C) Immunofluorescent staining of macrophages in the TA muscle, 3 days post CTX. Dual staining for Mac-3^+^ (red) and GFP^+^ (green) was done to identify differences in GFP^+^ macrophage recruitment between O-O and Y-O. No difference in macrophage recruitment was detected by after Mac-3 signal intensity was normalized to DAPI signal intensity. D) Flow cytometry gating of macrophages in naïve muscle and at 3- and 10-days post CTX injury (CD45^+^CD11b^+^F4/80^+^). E) Flow cytometry gating of Ly6C^hi^ and Ly6C^lo^ monocytes (Ly6C^+^Ly6G^−^) and neutrophils (Ly6C^+^Ly6G^+^). Percentages of different cell populations are shown in the indicated box. All data are presented as mean±SEM. All analyses were done using a 2-way ANOVA (≥2 factors). **P<0.05, **P<0.01, ***P<0.001 delineate differences within Y-Y or Y-O cohorts. #P<0.05 delineate differences between Y-O and O-O cohorts. BMT, Bone Marrow Transplant; TA, Tibialis Anterior; CTX, Cardiotoxin; Con, Saline Control.

We next assessed the influence the BM age has on the myogenic response (Figure 6). We first assessed how WT young or old muscles respond to injury and demonstrated that aging strongly influences myogenic capacity, in agreement with other studies (Figure 6A). We grouped myogenic genes based on their function: Pax7 is a satellite cell marker; Myogenin (MyoG) and MyoD are early myogenic markers representing commitment and differentiation of myoblasts; MCK, Myh3 and Myh4 are late myogenic markers, representing maturation of the myofiber. Myh3 is expressed in newly formed fibers and is gradually replaced by Myh4. Pax7 and MyoD expression was significantly higher in young muscle at 3- and 10-days post injury compared to old muscle. In young muscle, MyoG expression was elevated at 3 days post CTX and returned to baseline levels by 10 days post injury. Old muscle showed similar patterns of MyoG expression however this was significantly lower than in young muscle. In young muscle, late myogenic markers (MCK and Myh3) increased by 3 days post CTX while Myh4 expression was not upregulated until 10 days post injury. Compared to young muscle, MCK and Myh3 expression was significantly lower in old muscle at 3 days post CTX. This trend extended to 10 days post CTX when Myh4 expression was higher in young muscle.

**Figure 6.**
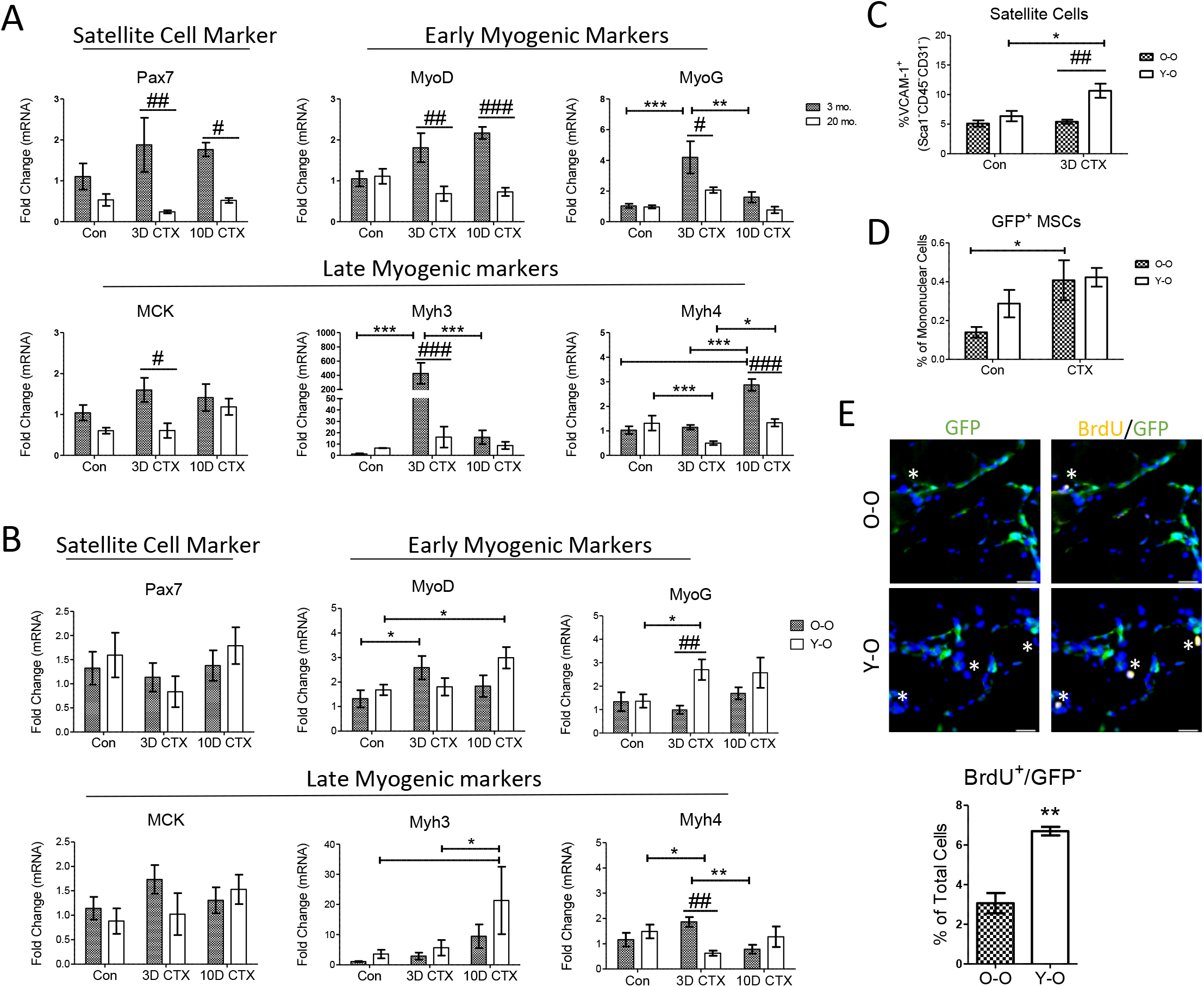
Bone marrow cell age influences myogenesis. A) Gene expression analysis of satellite cell and early and late myogenic markers in the TA muscle from WT young (3 month old) or old (20 month old) mice. The qRT-PCR data are reported as relative to housekeeping gene, Hprt (n=4). The control group is TA muscle that received an injection of saline. ***P<0.05, **P<0.01, ***P<0.001 delineate differences within young or old cohorts. #P<0.05 delineate differences between young and old cohorts. B) Gene expression analysis of satellite cell and early and late myogenic markers in the TA muscle of CTX-injured mice after heterochronic BMT. All recipients (hosts) were >18 months old at the time of reconstitution and underwent a 12-week recovery period before CTX administration. Mice that received young Sca-1^+^ BM cells are labelled Y-O. Mice that received old Sca-1^+^ BM cells are labelled O-O. The qRT-PCR data are reported as relative to housekeeping gene, Hprt (n=4). The control (Con) group is TA muscle that received an injection of saline. Con n=7, 3D CTX n=7, 10D CTX n=6. C-D) Flow cytometry analysis of GFP^−^satellite cells and GFP^+^ mesenchymal stem cells (n=4) and corresponding flow panels in uninjured or 3D post CTX injured muscle. C) Satellite cells are plotted as %VCAM-1^+^ of the Sca-1^−^, CD45^−^ and CD31^−^ population. D) Flow cytometry data of MSCs are plotted as % Sca-1^+^, CD45^−^ and CD31^−^ of mononuclear cells. Naïve flow cytometry data of SCs and MSCs is derived from Figure 3J-K. E) Young bone marrow induces host cell proliferation. BrdU^+^/GFP^−^ cells were quantified 3 days post CTX in Y-O and O-O mice. The top panels show representative images of staining for GFP (Green) and BrdU (Yellow) of TA cryosections of Y-O and O-O mice at 3 days post CTX. The bottom panel quantifies the number of BrdU^+^/GFP^−^ cells per total number of cells per image (n=3). All data are presented as mean±SEM. All analyses were done using an unpaired t-test (2 groups) or two-way ANOVA (≥2 factors). ***P<0.05, **P<0.01, ***P<0.001 delineate differences within Y-Y or Y-O cohorts. #P<0.05 delineate differences between Y-O and O-O cohorts. BMT, Bone Marrow Transplant; TA, Tibialis Anterior; CTX, Cardiotoxin; Con, Saline Control; MSC, Mesenchymal Stem Cell.

CTX injury on Y-O or O-O BMT mice resulted in some age-related differences that were partially rescued by young bone marrow. Notably, MyoG expression was higher at 3 days post CTX in Y-O, potentially representing differences in satellite cell proliferation (Figure 6B). In contrast, Myh4, a late myogenic marker, was downregulated at 3 days post CTX in the Y-O group. No significant changes were observed in Pax7 or MCK expression but MyoD and Myh3 expression trended higher in the Y-O cohort at 10 days post injury. Compared to control muscle, the satellite cell population increased at 3 days post CTX in Y-O but not O-O muscle (Figure 6C). There was no difference in GFP^+^ MSCs between Y-O and O-O after injury, though GFP^+^MSCs increased in the O-O group from control levels (Figure 6D). We next assessed whether BM aging had an impact on host cell proliferation. Three days post CTX, reconstituted mice were treated with BrdU to identify proliferating cells. In both Y-O and O-O mice we detected BrdU^+^/GFP^−^ cells (stars) and there were significantly more proliferating cells in mice that received young Sca-1^+^ cells (Figure 6E).

### M1 cytokines are an important part of muscle repair which becomes defective with age

One of the proposed mechanisms of muscle repair is that M1 macrophages and cytokines promote myoblast proliferation (and inhibit differentiation) while M2 macrophage promote myogenesis ^14,21^. Our BMT and aging data showed that Ly6C^hi^ monocytes and M1 cytokines (e.g. IL-1β and IFNγ) are part of the muscle repair process that is affected with age however this was done using gross muscle tissue. To correlate cytokine expression to immune cell type we analyzed microarray expression data from previously published data (GSE71152) ^22^ of Gr1^+^ monocytes and Cx3cr1^hi^ macrophages isolated from CTX injured muscle of 8-10 week old mice at 1-2-4- and 8-days post injury (Figure 7A). Both monocytes and macrophages expressed IL-1β early in muscle repair, from 1-2 days post injury while IFNγ was expressed at later stages from day 4-8 post injury. To determine age-related defects of expression of these cytokines we cultured peritoneal macrophages from young or old mice *in vitro* and evaluated cytokine expression after stimulation with LPS. IL-1β, IL-12a, IL-12b and IL-23 expression was consistently higher in young LPS-stimulated macrophages compared to old LPS-stimulated macrophages (Figure 7B). IFNγ and IL-10 expression showed the opposite trend and was more highly expressed in old LPS-stimulated macrophages. iNOS expression was significantly induced upon LPS treatment. Arg1 expression was reduced with LPS stimulation but expression was lower in unstimulated old macrophages compared to unstimulated young macrophages.

**Figure 7.**
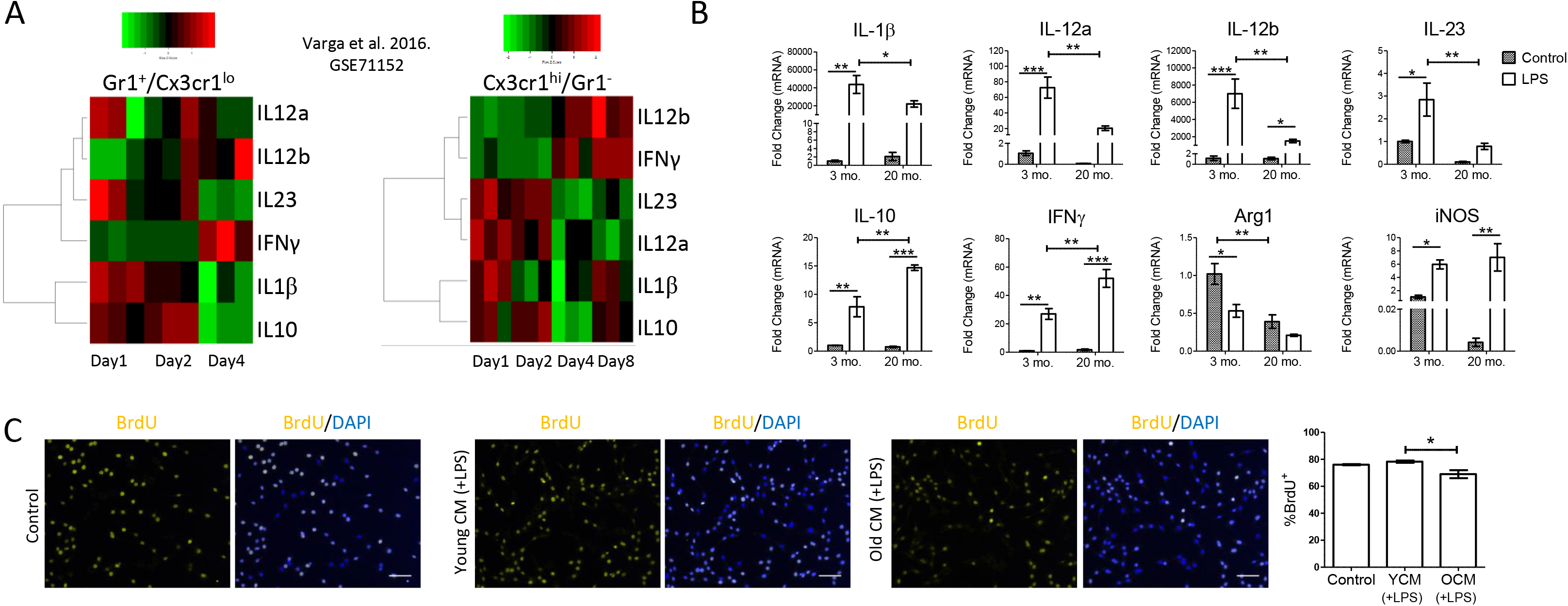
Age-related changes in M1 cytokines inhibits myoblast proliferation. A) Heatmap depictions of microarray gene expression values from Gr1^+^/Cx3cr1^lo^ or Cx3cr1^hi^/Gr1^−^ cells isolated from CTX injured muscle (GSE71152). Heatmaps were generates in RStudio. The timepoint of each analysis is indicated at the bottom of each heatmap (n=3). B) Peritoneal macrophages from 3- and 20-month old WT mice were cultured *in vitro* and stimulated using LPS (10ng/ml). Gene expression of Cytokines or functional M1 (iNOS) and M2 (Arg1) markers were evaluated. The qRT-PCR data are reported as relative to housekeeping gene, Hprt (n=3). C) Conditioned medium from old macrophages inhibits myoblast proliferation. C2C12 myoblasts were grown in conditioned medium isolated from young or old LPS-stimulated peritoneal macrophages for 24 hr. The degree of proliferation was quantified by BrdU^+^ nuclei (yellow). All analyses were done using a two-tailed unpaired t-test or a one- or two-way ANOVA (≥2 factors). Scale = 50 μm. *P<0.05, **P<0.01. Con, Control; LPS, Lipopolysaccharide; CM, Conditioned Medium.

To determine how macrophages affect myoblast proliferation, C2C12 myoblasts were cultured in conditioned medium (CM) collected from LPS stimulated young or old peritoneal macrophages. Cell proliferation was assessed by BrdU incorporation. Compared to serum-free control, CM from young LPS-stimulated macrophages had no effect on myoblast proliferation (Figure 7C). Interestingly, CM from old LPS-stimulated macrophages reduced cell proliferation. These *in vitro* studies demonstrate a mechanism whereby aging initiates a distinctly different immune response in terms of cytokine production that can have downstream consequences on the timing of myogenic repair processes such as myoblast proliferation. This finding also highlights the inhibitory effect of old immune cells on myogenesis.

## Discussion

The appreciation of inflammatory pathways as beneficial to tissue repair is increasing and harnessing the tools of macrophages, monocytes and T cells is bleeding into current and future treatment strategies. In skeletal muscle, the role of the inflammatory response in tissue repair has been investigated using various means of muscle damage in young mice, however the impact that aging has on this process is unclear. Recent studies have begun to dissect the impact that aging has on muscle repair by investigating intrinsic and extrinsic properties of satellite cells, the mediators of myogenesis. Here we show that CTX induces a rapid and trainset inflammatory response that is impaired with age and is associated with diminished myogenic capacity. Replacement of old BM with young BM restores the timing of the inflammatory cascade, and alters muscle repair in old mice via changes in muscle progenitor cells, demonstrating that extrinsic inflammatory cells affect satellite cell proliferation in old muscle. *In vitro,* myoblast proliferation was strongly affected by CM from old peritoneal macrophages, while CM from young peritoneal macrophages had no benefit. Together these data highlight that the old immune system is detrimental to myogenesis.

### Profiling the intersection between Inflammation and Aging on Muscle Repair

Recently Shavlakadze et al. profiled gene expression in the Gastroc, Liver, Kidney and Hippocampus across the lifespan of the rat from 6 to 27 months old and found that inflammation is a universal part of tissue aging ^23^. Our flow cytometry and gene expression profiles concur that aged muscles are deeply inflamed and provides a robust overview of the types of immune cells involved. In naïve muscle we observed more neutrophils and T cells and fewer satellite cells in old mice but higher levels of IL-1β, IFNγ and Il-12b. In contrast, after injury our gene and cell profiles demonstrate depressed/delayed expression of pro-inflammatory cytokines, fewer monocytes and delayed myogenesis in old muscle. This is in general agreement with a previous study that observed delayed inflammation and myogenesis in mice that received heterochronic autografts from young or geriatric mice ^18^. We did not, however, observe delayed myogenesis in old mice but a general impairment in early and late myogenic gene expression.

T cells, neutrophils, monocytes and macrophages are necessary for muscle repair and they are important regulators of myogenesis ^14,24^. Aging can therefore lead to two problems: i) Myogenic capacity is impaired (due to satellite cell niche instability and progressively, geriatric senescence ^7,8^. ii) A dysfunctional inflammatory response. As detailed in this study, local inflammation is high at rest and slow to activate in response to injury in old mice, a phenomenon observed by others ^25^. Therefore improving muscle repair processes in old mice will require tackling both satellite cell and immune cell deficiencies. To address one half of this dilemma we performed heterochronic BMT and found that young BM restores the timely recruitment of inflammatory cells and cytokines to injured muscle. Wang et al. recently performed heterochronic BMT and observed increased muscle mass by young BM cells in old mice while old BM had a negative impact on satellite cells in young mice ^19^. Although Y-O mice had an overall improvement in gross muscle mass and behavioural and locomotive tests, we did not see any change in satellite cell number or inflammation at baseline. Three days after CTX, however, mice that received young BM had more satellite cells and higher expression of MyoG. These mice also showed a progressive increase in Myh3 expression, at 10 days post CTX, indicating that muscle repair, though delayed, may be improved. Unfortunately, while inflammatory cell profiles were restored to trends observed in young WT mice, the myogenic response was not comparable, likely owing to the inherent aging defects associated with satellite cells.

### The M1 myeloid response in muscle repair

An important feature of muscle repair that has been well-documented is polarization of macrophages to M1 or M2 which inhibit and promote myogenesis, respectively ^24^. IL-10 is a classical M2 marker which can limit macrophage proliferation ^26^ and inflammatory cytokine production ^27^. Curiously at 3 days post injury IL-10 was upregulated in old but not young muscle and this trend was similar in LPS stimulated macrophages. This was initially surprising one may expect upregulation of an anti-inflammatory cytokine to be beneficial, and IL-10 is required for myogenesis post injury ^21^ and muscle-specific expression of IL-10 in aged muscle reduces inflammation and insulin resistance ^28^. However, IL-10 does not directly act on myoblasts but instead induces polarization of muscle-macrophages to M2, which then induce myogenesis via changes to the microenvironment ^21^. IL-10 can also suppress the M1 response by downregulation of M1 cytokines including IFNγ, IL-1β and IL-12b ^29–31^. With respect to aging, IL-10 expression is upregulated in aged muscle responsive for increased fibrosis via overactivation of M2a macrophages ^16^. Additionally, M2 macrophages increase while M1 macrophages decrease with aging in human muscles ^32^, demonstrating that restoration of the M1 response may be key to maintenance of muscle repair with age.

We examined expression of M1 cytokines IL-1*β*, IL-12b and IFN*γ* which are tightly aligned with the timing of the Th1 helper T cell response ^33^. During muscle repair in young and old WT mice, we observed changes in the IL-12-IFNγ signaling axis, which encompasses monocytes, macrophages, T cells and myoblast behaviour. IFNγ can stimulate M1 macrophage activity, including upregulation of IL-12b, which in turn may stimulate production of IFNγ in T cells, driving the feedforward loop ^33^. In muscle, IFNγ acts directly on muscle cells to repress myogenesis ^34^. At 3 days post injury, IFNγ was upregulated in Y-O but not O-O mice which could be an important step in myogenesis, that is to promote proliferation and repress differentiation. Using *mdx* mice to model muscular dystrophy, Madaro et al. demonstrated that loss of macrophages led to precocious differentiation of satellite cell, reduced self renewal capacity and induced commitment to an adipogenic lineage ^35^. In a similar vein, changing age-dependent changes in macrophage populations (or other immune cells) would impact cytokines such as anti-differentiation factor, IFNγ, and alter the timing between proliferation and differentiation. Several questions remain unanswered. Does inflammation in old muscle preclude the timely response to acute injury? Is it the preconditioned inflammatory state of old muscle, the impairs the mobilization/differentiation/activity of old inflammatory cells, or a combination of both that alters myogenesis? A careful understanding of the chronic vs acute actions of each cytokine is necessary for better muscle repair in older organisms.

## Conclusion

Aging promotes an inflammatory microenvironment in muscle, yet in response to acute injury the inflammatory response is dampened. Replacement of old BM with young BM restored the timely infiltration of myeloid cells to injured muscle and altered the myogenic response. These data demonstrate the importance of BM aging (extrinsic factors) on the microenvironment of aged skeletal muscle in satellite cell activation and differentiation, however they also demonstrate the intrinsic impairment that aged satellite cells possess. To produce a robust myogenic effect in the elderly, therapies should target both the inflammatory response and inherent defects in satellite cell pool.

## SMethods

### Bone marrow reconstitution

The Animal Care Committee of the University Health Network approved all experimental procedures which were carried out according to the Guide for the Care and Use of Laboratory Animals (NIH, revised 2011). Bone marrow from young (3 month) or old (18-20 month) C57BL/6-Tg(CAG-EGFP)1Osb/J enhanced GFP mice was isolated from the tibiae and femur as follows: Cells were dissociated from bone marrow in PBS using an 18G-23G needle. Cells were then incubated in 5 ml of Red Blood Cell lysis buffer (154.42 mM NH_4_Cl, 11.9 mM NaHCO_3_, 0.026 mM EDTA) for 5 min and then centrifuged for 5 min at 1000 rpm. These steps were repeated for a total of two times each. The cell pellet was suspended in PBS and passed through a 40 μm filter to remove debris. Sca-1^+^ cells were purified using Sca-1 magnetic purification (Stem Cell Technology). Old (>18-20 month) C57BL/6 female mice were lethally irradiated at 9.5 Gy and immediately received an infusion (through the tail vein) of 2×10^6^ GFP^+^/Sca-1^+^ cells from young or old GFP mice. Three months later, mice were sacrificed or underwent CTX muscle injury.

### Muscle injury

Cardiotoxin was administered as previously described ^36^. Briefly, mice were sedated using isofluorane and the hindlimbs were cleaned using ethanol. One, two, or three doses of 20 μl of 10 μM Cardiotoxin (Latoxan, L8102) was injected into the Tibialis anterior (TA), Quadriceps (Quad), or Gastrocnemius (Gastroc) muscles, respectively. The uninjured leg received a volume control injection of Saline into the TA muscle for qRT-PCR analysis. For CTX-BrdU experiments, CTX was administered as described above on day zero. Three days after CTX, 5-bromo-2′-deoxyuridine (BrdU; Sigma) was administered via intraperitoneal injection six hours prior to sacrifice at a dose of 50 mg/kg.

### Locomotive and behavioural tests

Open field test: The open field test apparatus consisted of a 38 x 60 x 60 cm chamber with grey Plexiglas walls and transparent ceiling to allow for video recording. Mice were placed in the chamber for 10 min and their ambulatory distance and rearing count was tracked and recorded. Video files were initially analyzed by idTracker ^37^ in order to obtain x- and y-coordinates associated to activity, followed by subsequent analysis using a custom python script for extraction and calculation of ambulatory distance and anxiety related measurements. The testing apparatus was cleaned with 70% ethanol between each mouse. Rotarod test: The accelerating rotarod test was performed on a Pan Lab Lsi Rota-Rod/RS Model 8200 and the times spent before falling were recorded. Prior to testing, the mice were acclimated to the device at a constant speed (4 rpm) and at constant acceleration (4 rpm to 40 rpm in 30 seconds) for three separate trials for each setting with at least a 10 min interval between each trial. Afterwards, mice were tested on the accelerating rotarod over the course of three separate trials that were averaged to produce the mean amount of time spent on the device before falling. The testing apparatus was cleaned with 70% ethanol between each mouse.

### Flow cytometry

Mononuclear cells were isolated from hindlimb muscle for flow cytometry as described ^38^. For analysis of uninjured muscle, all hindlimb muscle was collected from both legs after perfusion with PBS. Three or 10 days post CTX, cells were isolated from only the injured Gastroc and Quad muscles. Isolated mononuclear cells were stained at 4°C in FACS buffer (2% FBS in PBS). Antibodies used to detect satellite and mesenchymal stem cells: Sca-1 (Clone E13-161.7, Cat. no. 553336), Biotin anti-mouse CD106 (Clone 429 MVCAM.A, Cat. no. 105703) or Pacific Blue™ anti-mouse CD106 (Clone 429 MVCAM.A, Cat. no. 105722), CD45 (Clone 30-F11, Cat. no. 103112), CD31 (Clone MEC13.3, Cat. no. 102509), and Streptavidin BV-711 (Cat.. no. 405241). Antibodies used to detect Immune Cells: F4/80 (Clone BM8, Cat. no. 123113), Ly6C (Clone HK1.4, Cat. no. 128015), Ly6G (Clone 1A8, Cat. no. 127613), Cd11b (Clone M1/70, Cat. no. 101207), CD45 (Clone 30-F11, Cat. no. 103128 (Alexa Fluor 700) or 103108 (FITC), depending on if samples included GFP signal), CD3 (Clone 17A2, Cat. No. 100205) and CD4 (Clone GK1.5, Cat. No 100413). Antibodies were purchased from BioLegend and BD Biosciences. Neutrophils were identified as CD45^+^ CD11b^+^ F480^−^ Ly6C^+^ Ly6G^+^, monocytes were identified as CD45^+^ CD11b^+^ F480^−^ Ly6C (hi or lo) Ly6G^−^, macrophages were identified as CD45^+^ CD11b^+^ F480^+^, and T cells were identified as CD45^+^ CD3^+^ CD4^+^. Data was acquired on an LSR II (BD Biosciences) flow cytometer and data analyzed using FlowJo software. Representative images of gating strategies are shown in Supplemental Figures S2-S5.

### qRT-PCR

Tissue was snap frozen in liquid nitrogen and then ground into a fine powder. RNA was isolated using TRI-Reagent (Sigma) according to the manufacturer’s instructions. cDNA was prepared from 1000ng of RNA using NxGen M-MulV Reverse Transcriptase (Lucigen; 30222-1) according to the manufacturer’s instructions. cDNA expression was analyzed using SensiFAST (Bioline-98005) SybrGreen using the following parameters: 95° 2 min; [95° 5 s; 60° 10 s; 72° 20 s for 40 cycles]. Relative quantification of gene expression was calculated by ∆∆CT method with Hprt or β-actin as housekeeping genes, where indicated. Experiments were based on an average of two technical replicates. The number of biological replicates used are indicated in the figure legends. Primers are listed in Supplemental Table S1.

### Immunofluorescence and histology

The TA muscle was placed in 2% PFA and incubated in 10%, 20% and finally 30% Sucrose, overnight. Muscle tissue was embedded in OCT and sectioned at 7 μM. 2% PFA was added to tissue sections for 10 minutes and washed 3X with 1% BSA in PBS. For BrdU analysis, tissue sections were placed in 2M HCl at 37°C for 30 minutes and washed 3X with 1% BSA-PBS. Next, the tissue was incubated with borate buffer (0.1M Boric acid, pH 8.5) at room temperature for 10 minutes and washed 3X with 1% BSA-PBS. Tissue was permeabilized in 0.5% Triton-X and blocked in 10% BSA for 15 minutes. Antibodies are as follows: Primary antibodies: BrdU (Abcam, ab6326), CD45 (BD, 550539), GFP (ThermoFisher, A21311) and Laminin (Sigma-Aldrich, L9393). Secondary antibodies: Goat anti-Rat Alexa 546 (ThermoFisher, A11081), Chicken anti-Rat, Alexa 647 (ThermoFisher, A-21472) and Donkey-anti-Rabbit (A31573). with 4′,6-diamidino-2-phenylindole (DAPI, Sigma-Aldrich) was used to counterstain nuclei. BrdU^+^/GFP^−^ nuclei were quantified using an average of three fields of view within the injured TA, followed by the average of BrdU^+^/GFP^−^ cells over three independent experiments (biological replicates). For histological analysis the TA muscle was placed in formalin for 24 hr, washed in 70% ethanol and embedded in paraffin wax. Muscles were sectioned at 7 μm. Hematoxylin and Eosin (H&E) staining was completed by the pathology lab at University Health Network (Toronto ON). The sections were viewed and photographed using the Olympus Slide Scanner CM-10 or the Zeiss LSM700 confocal microscope, and the digital images were processed with Zeiss ZEN. The cross-sectional area of 60-100 myofibers per sample was calculated using ImageJ software (n=5).

### Isolation and in vitro culture of peritoneal macrophages

Young (3 month) or old (18-20 month) mice were given an intraperitoneal injection with 4% thioglycolate three days before sacrifice. On the third day, mice were sacrificed via carbon dioxide inhalation followed by cervical dislocation. Cells were collected by peritoneal lavage and were resuspended in DMEM/F12 media (ThermoFisher) supplemented with 10% FBS and 1% Penicillin-Streptomycin (P/S). Cells were seeded in 12-well plates at 10^6^ cells/ml. The cells were allowed to adhere overnight, and the media was changed the next day before experimentation. LPS was added to cells at a concentration of 10 ng/ml for 4 hr before collection of RNA.

### C2C12 culture experiments

C2C12 myoblasts (ATCC) were seeded into a 24-well dish at 2×10^4^ cells/ml in DMEM (Gibco) supplemented with 10% FBS and 1% P/S. The next day, conditioned medium from young or old peritoneal macrophages were added to culture. Conditioned medium was collected as follows: Cells were seeded into 6-well plates at 10^6^ cells/ml and stimulated with LPS as described above. After macrophage activation, medium was washed twice with PBS and replaced with 1.5 ml serum-free DMEM/F12 media for 24 hr. Conditioned media was collected, supplemented with 10 μM BrdU, and added to proliferating C2C12 myoblasts one day after seeding. Control cells received serum-free DMEM+1% P/S. After 24 hrs, C2C12 myoblasts were fixed in 2% PFA and prepared for immunofluorescent staining as described above. BrdU^+^ nuclei were quantified on ImageJ and normalized to the total number of nuclei. Nuclei were averaged from two fields per biological replicate (n=3).

### Statistics

Data are depicted as mean ± SEM. Experiments with two groups were analyzed using a two-tailed unpaired t-test. Experiments with one variable and multiple groups were analyzed using a one-way ANOVA followed by Tukey’s post hoc test; Experiments with two variables were analyzed using a two-way ANOVA followed by Bonferroni’s post hoc test (GraphPad Prism). Significance was accepted as P < 0.05.

## Author Contributions

S.W.T. designed the experiments and wrote the manuscript. Bone marrow reconstitution experiments were completed by S.W.T, F.A., L.W., A.Y., and J.W. S.W.T and A.Y. contributed to immunofluorescence experiments. L.W. and S.M. performed and analyzed open field and rotarod experiments. S.W.T. and F.A. contributed to flow cytometry experiments. S.W.T completed gene expression analysis, C2C12 myoblast experiments and CTX treatments. F.A. completed *in vitro* experiments involving peritoneal macrophages. F.A., and R-K. L. edited the manuscript.

## Supplemental Information

**Supplemental Table 1.**
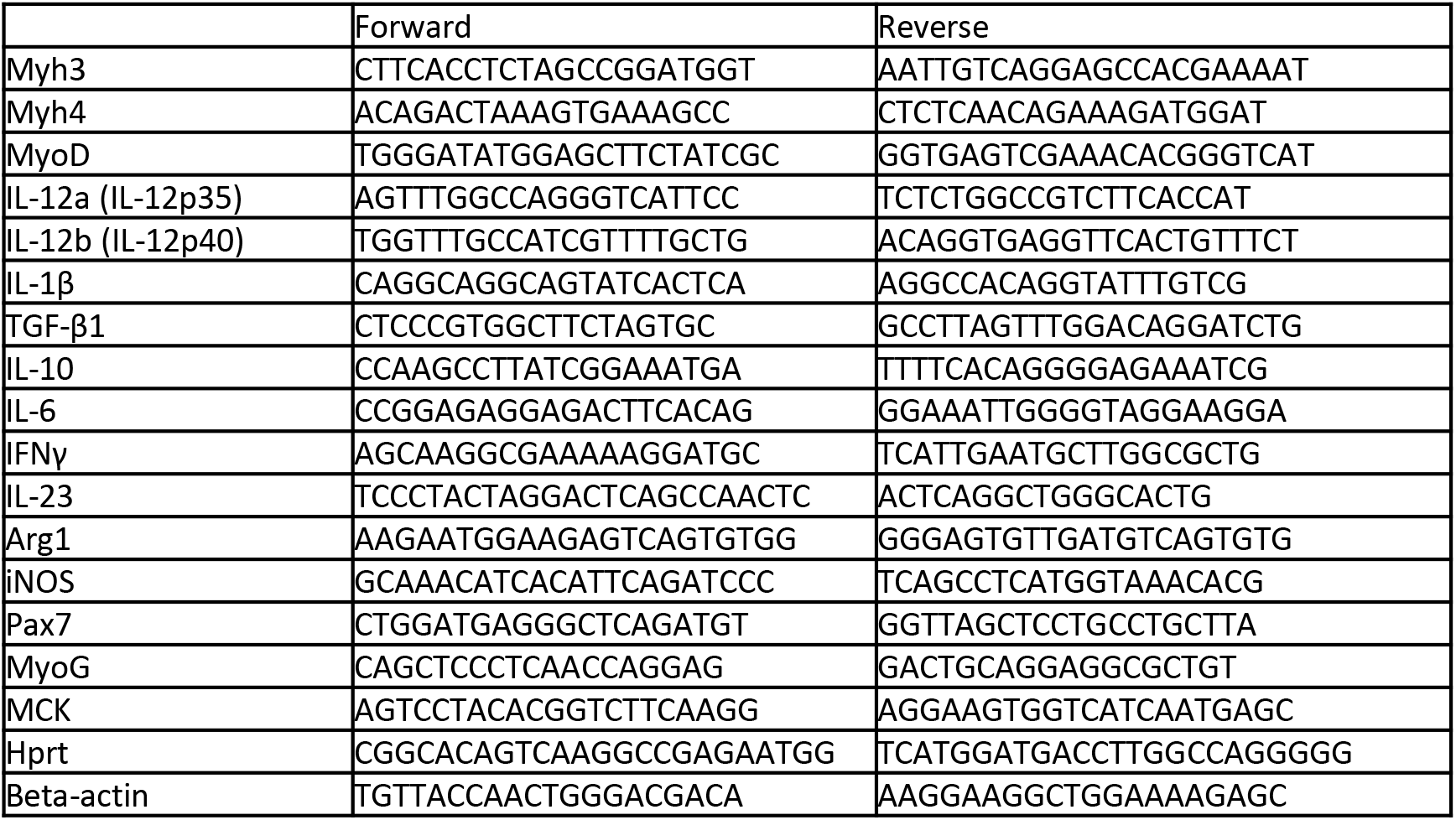
Primers used for qRT-PCR (mouse).

**Supplemental Figure S1.**
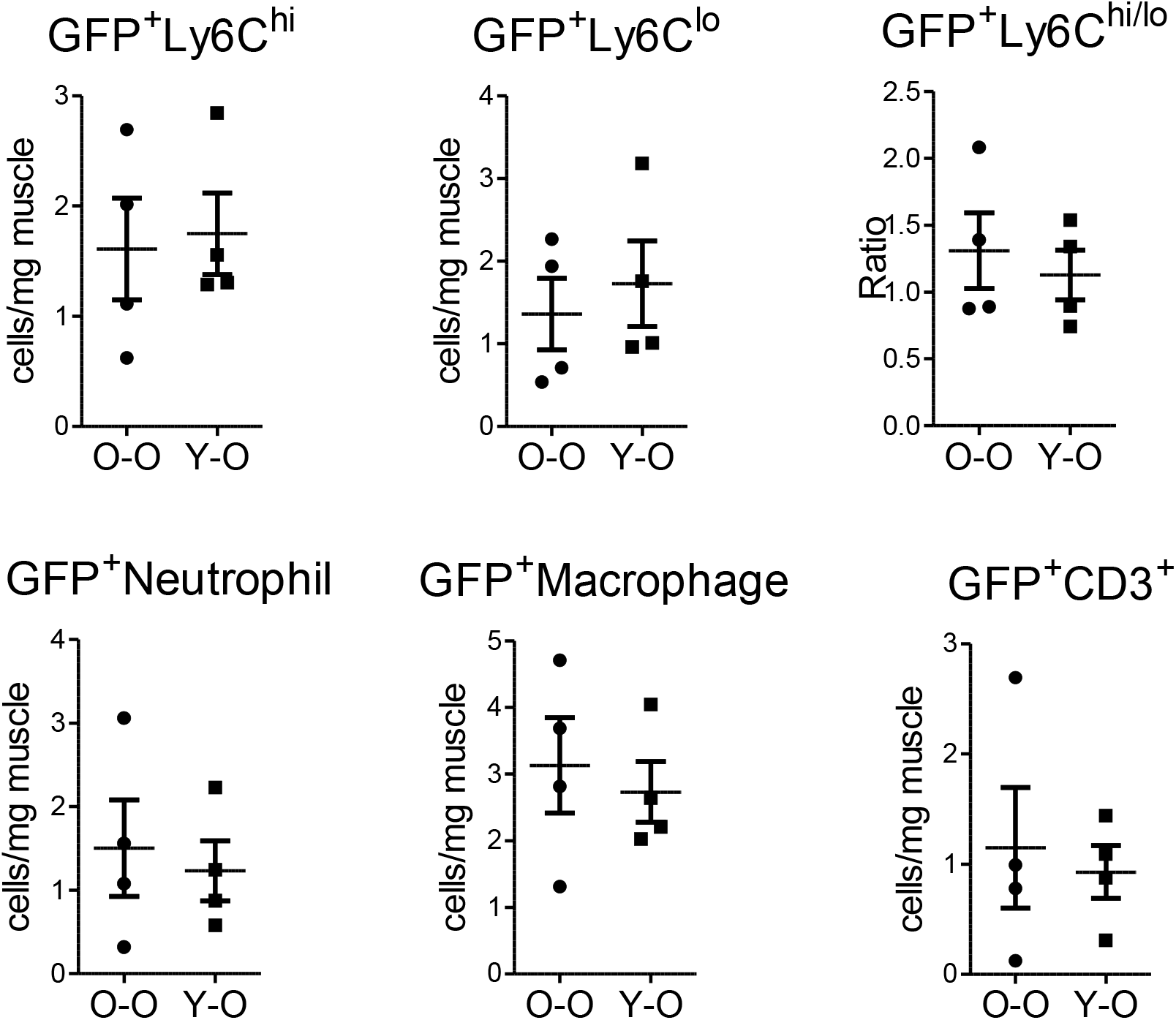
Identification of immune cells by flow cytometry after heterochronic BMT.

**Supplemental Figure S2.**
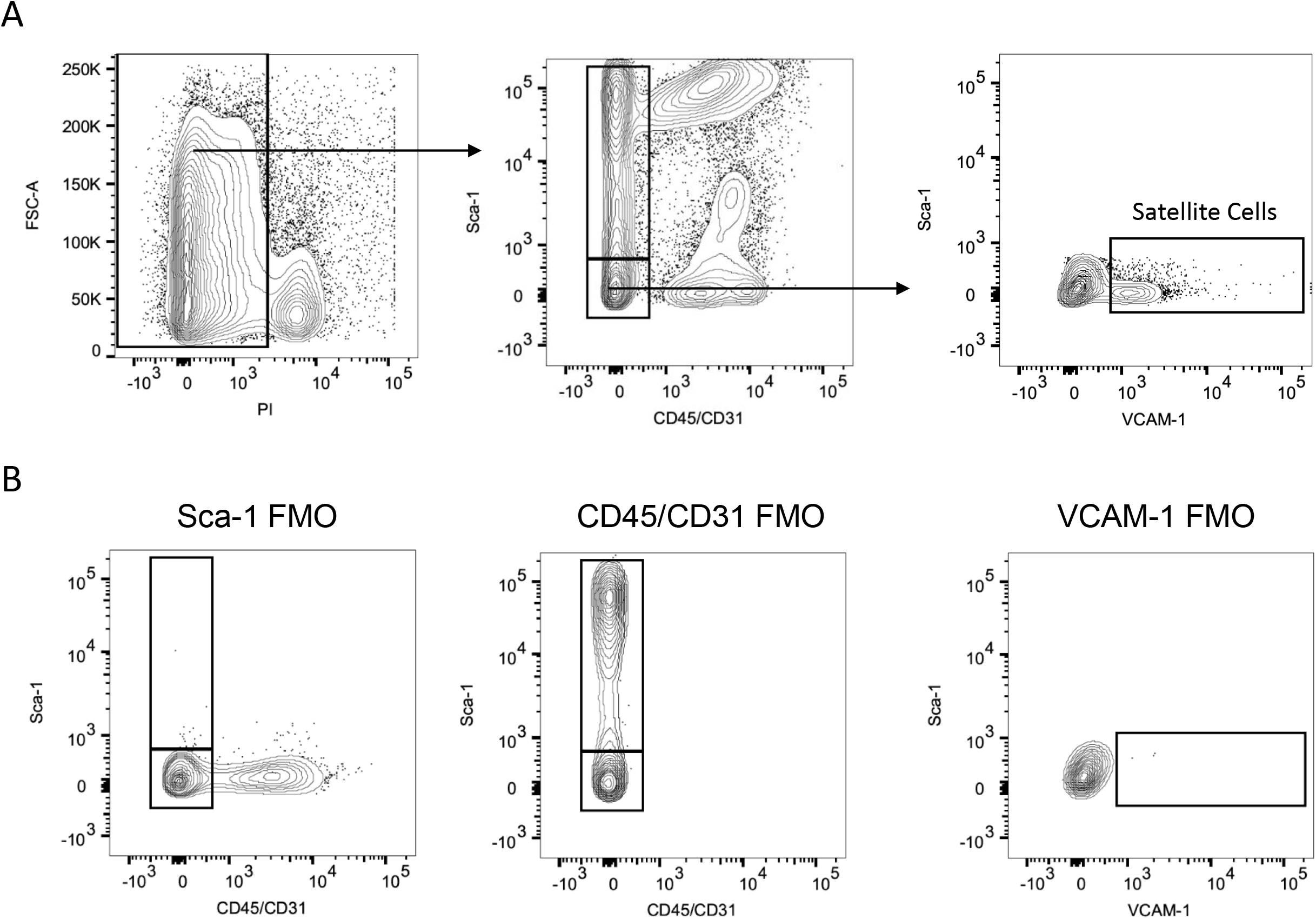
Identification of satellite cells by flow cytometry. A) Representative image depicts isolation of satellite cells in a young, WT naïve mouse. B) Representative images of (Fluorescence Minus One) FMO controls.

**Supplemental Figure S3.**
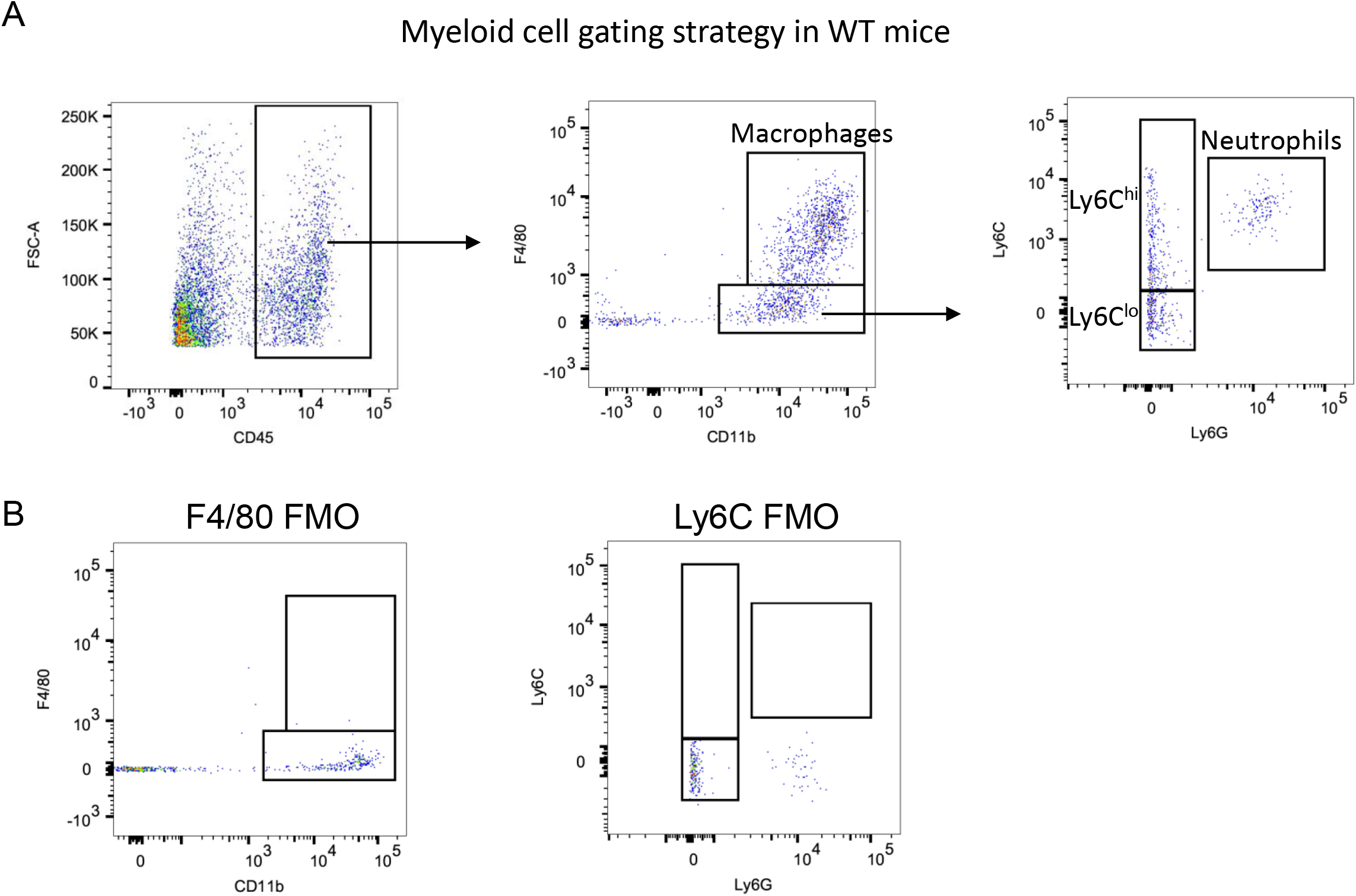
Identification of immune cells in WT mice by flow cytometry. A) Gating strategies to identify CD45 cells, macrophages, neutrophils and monocytes. B) Representative images of (Fluorescence Minus One) FMO controls.

**Supplemental Figure S4.**
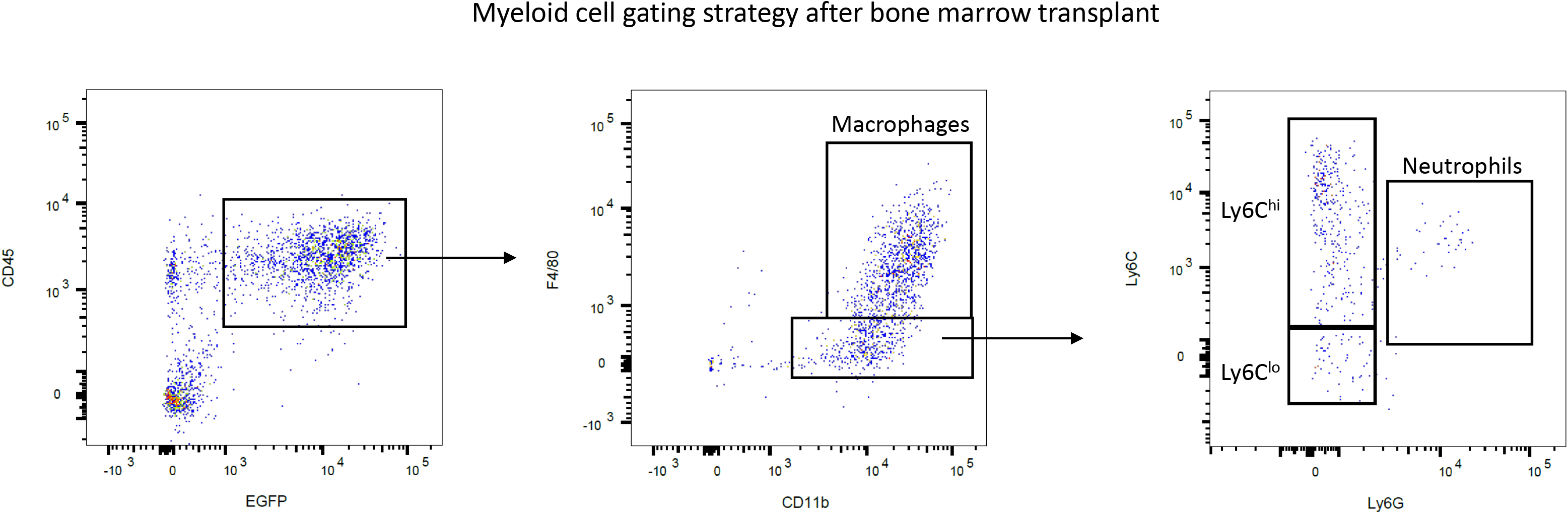
Identification of immune cells in mice after bone marrow transplant of GFP^+^Sca-1^+^ cells by flow cytometry. Gating strategies to identify CD45 cells, macrophages, neutrophils and monocytes.

**Supplemental Figure S5.**
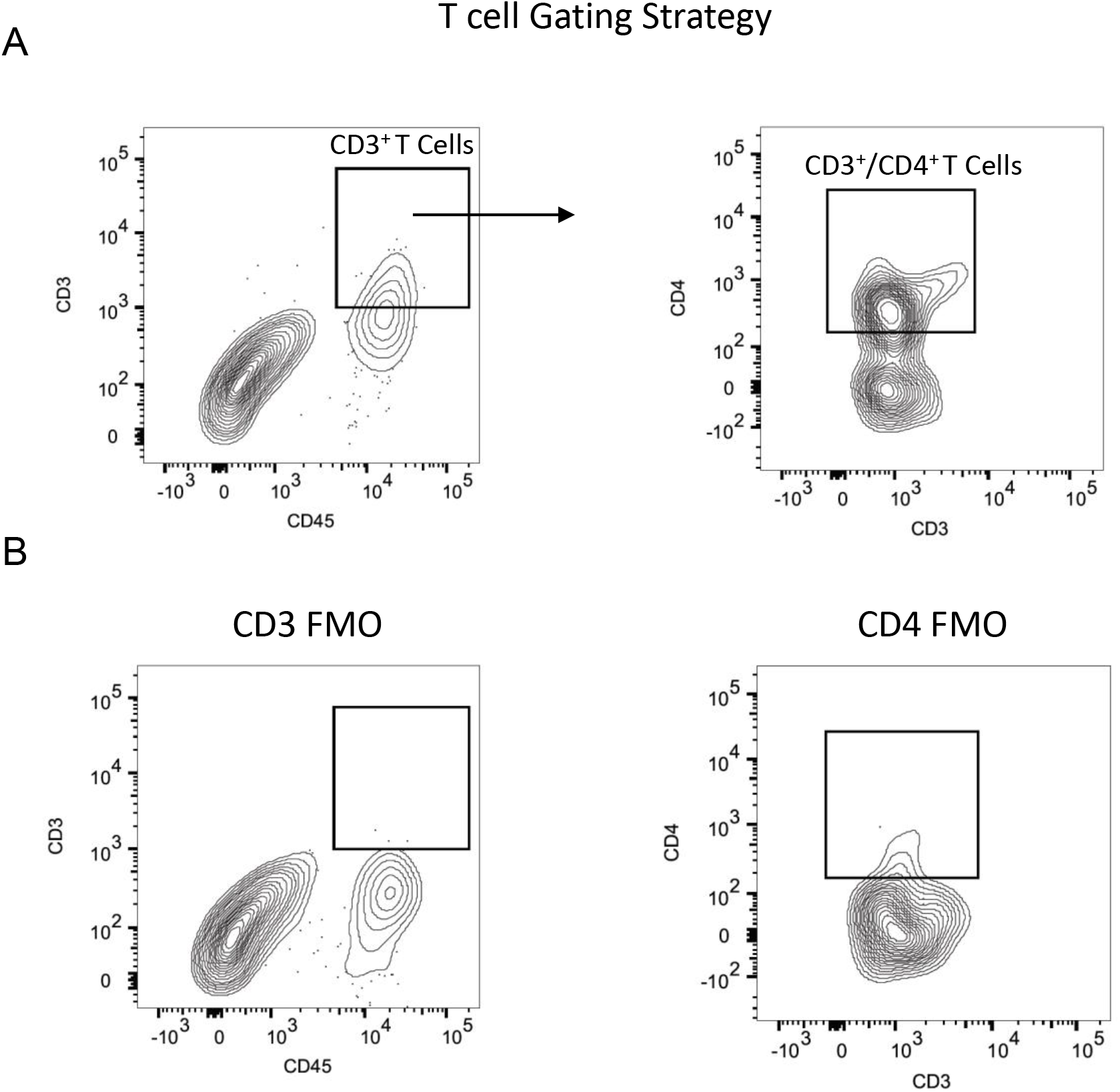
Identification of T cells in WT mice by flow cytometry. A) Gating strategy to identify CD45^+^/CD3^+^/CD4^+^ T cells B) Representative images of (Fluorescence Minus One) FMO controls.

